## Supplementary figures and images for "Effects of cellular stress on pigment production in *Rhodotorula mucilaginosa/alborubescens* AJB01 strain from the Caribbean region of Colombia"

### Figure S1

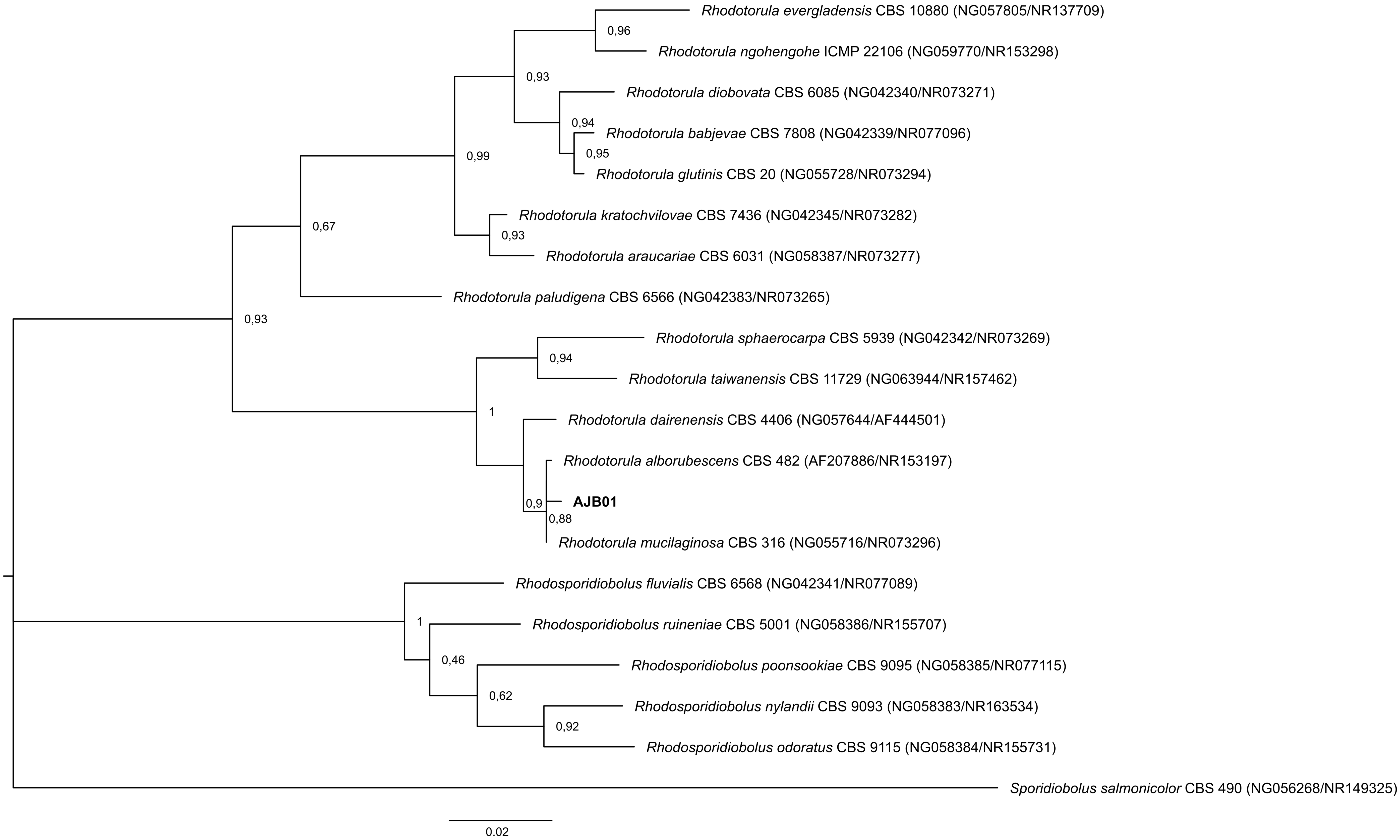
